# Diachronic monitoring of snow leopards at Sarychat-Ertash State Reserve (Kyrgyzstan) through scat genotyping: a pilot study

**DOI:** 10.1101/835025

**Authors:** Julie Rode, Agnès Pelletier, Julien Fumey, Stéphane Rode, Anne-Lise Cabanat, Anne Ouvrard, Bastien Chaix, Bradley White, Matthew Harnden, Nguyen Thi Xuan, Alexander Vereshagin, Didier Casane

**Affiliations:** Objectif Sciences International NGO, OSI-Panthera program, Geneva, Switzerland; WildMap Consulting Ltd., Fort St. John, British Columbia, Canada; Human Genetics and Cognitive Functions, Institut Pasteur, UMR3571 CNRS, Université de Paris, Paris 75015, France; Natural Resources DNA Profiling & Forensic Centre, Peterborough, Ontario, Canada; Sarychat-Ertash State Reserve, Kyrgyzstan; Issyk-kul Biosphere Reserve, Kyrgyzstan; Évolution, Génomes, Comportement, Écologie. CNRS, IRD, Université Paris-Sud. Université Paris-Saclay. F-91198 Gif-sur-Yvette, France; Université Paris Diderot, Sorbonne Paris Cité, France

**Author notes:** These authors equally contributed to this work (Julie Rode, Agnès Pelletier).

**Keywords:** snow leopard, noninvasive genotyping, population dynamics, microsatellite, relatedness, diachronic monitoring, citizen science, Central Asia

## Abstract

Snow leopards (*Panthera uncia*) are a keystone species of Central Asia’s high mountain ecosystem. The species is listed as vulnerable and is elusive, preventing accurate population assessments that could inform conservation actions. Non-invasive genetic monitoring conducted by citizen scientists offers avenues to provide key data on this species that would otherwise be inaccessible. From 2011 to 2015, OSI-Panthera citizen science expeditions tracked signs of presence of snow leopards along transects in the main valleys and crests of the Sarychat-Ertash State Reserve (Kyrgyzstan). Scat samples were genotyped at seven autosomal microsatellite loci and at a X/Y locus for sex identification, which allowed estimating a minimum of 11 individuals present in the reserve from 2011 to 2015. The genetic recapture of 7 of these individuals enabled diachronic monitoring, providing indications of individuals’ movements throughout the reserve. We found putative family relationships between several individuals. Our results demonstrate the potential of this citizen science program to get a precise description of a snow leopard population through time.

## Introduction

The most important threats facing snow leopard (*Panthera uncia*) populations include habitat loss, loss of prey-resources, human-wildlife conflicts, and poaching ^1–4^. Because of these, snow leopards were listed as Endangered by the IUCN from 1986 to 2016. Thanks to recent conservation efforts, the species was downlisted to Vulnerable (C1) in 2017. Rough estimates of the total number of individuals indicate between 2,700 and 3,400 snow leopards across the species range (https://www.iucnredlist.org/species/22732/50664030). However, many concerned countries are missing up-to-date information regarding snow leopard population sizes and demographic trends because they are highly elusive carnivores who live in remote mountainous locations, making ecological, behavioural and population studies challenging ^4,5^. Research that seeks to provide current and accurate demographic trends is the main priority highlighted in the Snow Leopard Survival Strategy ^2^.

In Kyrgyzstan, snow leopard numbers have decreased at an alarming rate over the last few decades, with 650-800 individuals estimated in the 1990s, against 150 – 200 in 2000 ^6^. Latest estimates are around 350 – 400 individuals (National Academy of Sciences of Kyrgyzstan, unpublished data) ^4^. So far, protection efforts in this country have mainly focused on preventing poaching, one of the most important threats to wildlife since the break-up of the Soviet Union ^6^. The main protected area, the Sarychat-Ertash State Reserve, established in 1995, is located in the Tien-Shan mountain range of Kyrgyzstan (Fig. 1). Beside poaching, major threats on biodiversity in the Sarychat-Ertash State Reserve (SESR) include climate change, mining, overgrazing, and overhunting ^7^. The SESR highlands are also surrounded by areas where hunting is authorized, which increases pressure on snow leopards and their prey. The SESR is divided into fourteen districts, each monitored by a ranger. Several studies based on genetic analyses and camera trapping estimated the snow leopard population size in the SESR to be around 20 individuals in 2011 ^4,8,9^. However, a long term accurate picture of population dynamics is lacking. Non-invasive genetic analyses are suitable tools to provide estimates of demographic parameters necessary to frame conservation plans ^10–13^.

**Figure 1.**
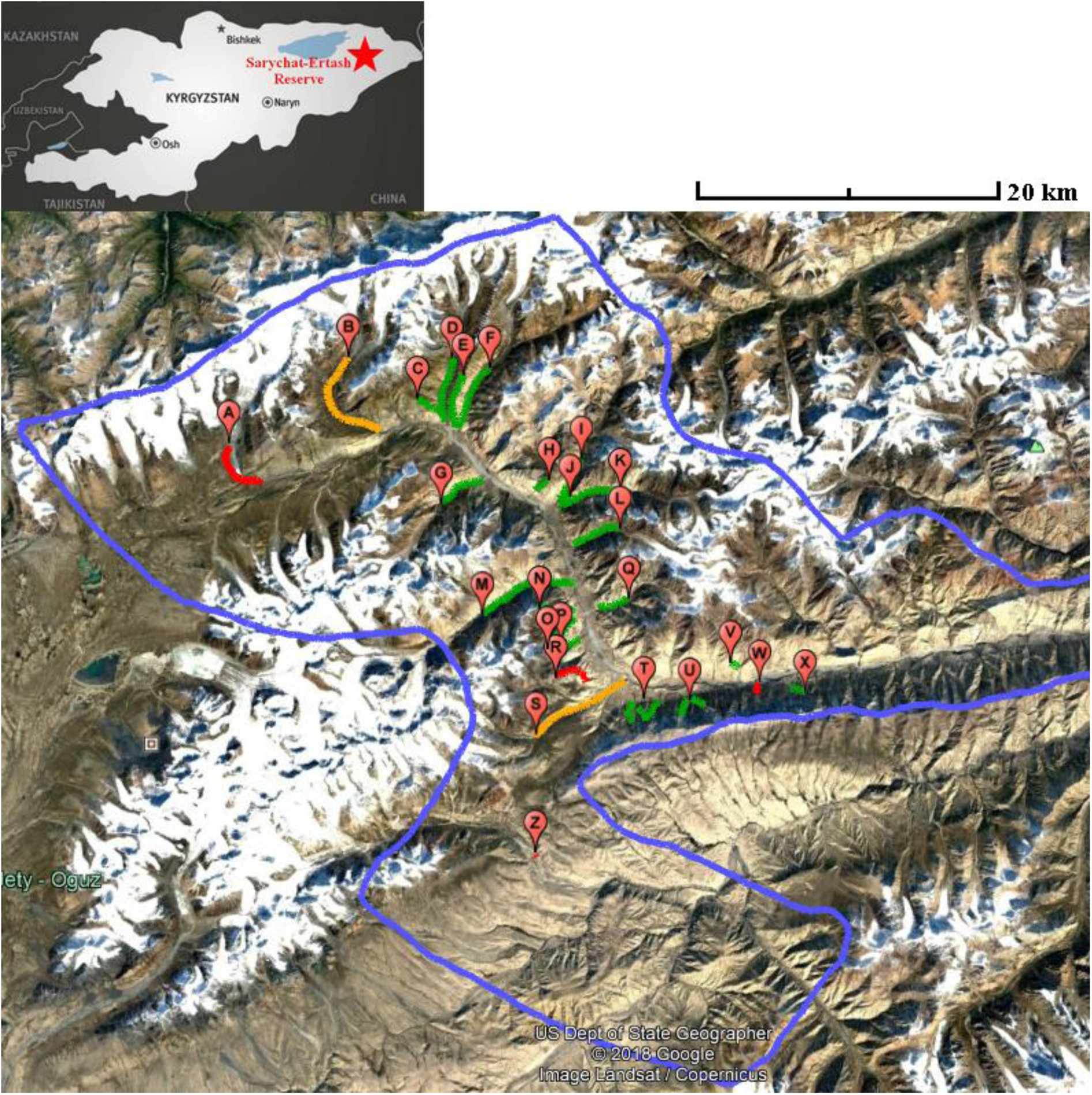
Sampling area. A) Sarychat-Ertash State Reserve location in Kyrgyzstan; B) snow leopard signs and sampled scats along transects within SESR from 2011 to 2015. Blue line: SESR boundary; red line: transects where no snow leopard signs were found; orange line: transects where snow leopard signs was found; green line: transects where snow leopard scats were sampled. Capital letters refer to transect indexes which are compiled in table S1 in supplementary materials. The framed area is shown in Fig.2.

The main objectives of the present study were to assess the suitability of non-invasive genetic techniques to study the snow leopard population living in the SESR, and to obtain information on family relationships and movement patterns. All samples obtained in the field were collected through seasonal citizen science expeditions where participants were trained by local rangers and scientific experts on the methodology of scat sampling. Despite the practical constraints associated with this kind of study for such an elusive species, we were able to identify several individuals, follow their movements over several years, and find evidence of relatedness between some of them.

## Materials and methods

### Study Area

Before 1995, the SESR was a grazing area for USSR (Union of Soviet Socialist Republics) shepherds who lived there year long with flocks containing thousands of animals. The SESR boundary encompasses 1,340 km², with a 720 km² core zone and a 620 km² buffer zone ^7^ (Fig. 1). The relief is characterized by large flat valleys about 1 km wide surrounded by high mountains of altitudes ranging from 2,000 to 5,500 m ^7^. The climate is continental, with low average temperatures even during the summer months (−21.5 °C in January; +4.5 °C in June). Vegetation types include arid grasslands and alpine meadows, with a majority of bushy and blanket cover type plants that are able to sustain the harsh and windy climate ^7^. Beside the snow leopard, several large and mesocarnivores are found in the reserve: wolves (*Canis lupus*), red foxes (*Vulpes vulpes*), Tian Shan brown bears (*Ursus arctos isabellinus*), manuls (*Felis manul*), Eurasian lynx (*Lynx lynx isabellinus*), and several mustelids. Large and medium herbivores, which are snow leopards’ usual prey species, are also found, including Siberian ibex (*Capra sibirica*), argali (*Ovis ammon*), and grey marmots (*Marmota baibacina*). Several species of birds are present, a few of which represent prey species for snow leopards, such as snowcocks (*Tetraogallus himalayensis*) and chukar partridges (*Alectoris chukar*) ^4,7^.

### Monitoring of snow leopard presence

In this study, snow leopards of SESR were monitored during citizen science expeditions led by the OSI-Panthera research program (osi-panthera.org). These 2 to 4 weeks expeditions were conducted by local rangers and guides, scientific educators, and volunteers. Monitoring effort increased over time, with 2 expeditions in 2011 (July and August), 3 in 2012 and 2013 (June, July, and August), and 4 in 2014 and 2015 (June, early July, July-August and late August).

Snow leopard presence was recorded based on specific signs (presence of scats, hairs, scratch marks, tracks, urine sprays on rocks, and carcasses of prey species), and based on pictures from camera traps set at known locations. Incidental species were also recorded to obtain information on prey presence and biodiversity level.

The protocol consisted of prospecting for snow leopards signs along transects (Fig. 1). As snow leopards are more likely found in steep and rocky environments and travel along topographic edges ^4^, transects were designed along waterbodies, ridgelines and cliffs, as well as in narrow valleys and canyons ^9^. Most transects were set within a sampling area of about 500 km² within the SESR core zone, around the main valley in which the Ertash River flows, and at the entry of secondary valleys (Fig. 1).

Glaciers, which are not considered high quality habitat for snow leopards ^4^ were only prospected once due to both low accessibility and time constraints. As snow leopards are territorial and travel several kilometers each day ^4^, our large sampling area enabled us to estimate the number of individuals in the whole reserve and to assess their movements. The list of prospected ridgelines and information on the presence of putative snow leopards signs and collected feces can be found in **Table S1**.

### Collection of scat and hair samples

Putative snow leopard scats were identified based on size, shape, vegetation content, as well as proximity to tracks, scratch marks and carcasses. Scats were collected with latex gloves, then stored in silica beads containers at room temperature until DNA extraction. A total of 137 putative snow leopard scat samples were collected and subsequently analyzed (14 in 2011, 24 in 2012, 45 in 2013, 41 in 2014, and 13 in 2015). Hair samples from 2 captive snow leopards (1 male and 1 female) with known pedigrees were collected by the staff of Jungle Cat World Wildlife Park (Ontario, Canada), and were used as positive controls for species identification and sexing. Additional scat and hair samples from other captive snow leopards were collected by the staff of the Toronto Zoo (Ontario, Canada) and were also used as positive controls during genetic analyses.

### DNA extraction

DNA from both hair and scat samples was extracted with a Qiagen DNeasy Blood and Tissue extraction kit. Depending on the sample, 10 hairs with visible roots or the outer mucosal layer of dried scat (obtained by swabbing scats with water hydrated cotton-tipped applicators) were used for extraction. These were suspended in 180 μl of lysis buffer (2 M urea, 0.1 M NaCl, 0.25% N-laurylsarcosine, 5 mM CDTA and 0.05 M Tris HCl pH 8). For scat, protein digestion was performed by adding 20 μl of proteinase K (600 U/mL) to each sample, and by incubating the samples in a 65 °C water bath for 1.5 hours, followed by a second spike of proteinase K, and overnight incubation at 37 °C. For hair, protein digestion was conducted by adding 25 μl of proteinase K (600 U/mL) and 10 μl of 1M DTT to each sample, and by incubating the samples in a 65 °C water bath for 1.5 hours. This was followed by a second spike of proteinase K and DTT, and overnight incubation at 37 °C. Subsequent extraction steps were carried out as per the manufacturer’s instructions, following the protocol with final DNA elution to 50 μl TE. Extracted DNA concentrations of the hair samples were measured using the Quant-IT PicoGreen dsDNA kit (Invitrogen) with a BMG FLUOstar fluorometer following the manufacturer’s protocol.

### Species identification

Snow leopard identification through scat was carried out using two snow leopard specific primers, CYTB-SCT-PUN-F’ (5’-TGGCTGAATTATCCGATACC) and CYTB-SCT-PUN-R’ (5’-AGCCATGACTGCGAGCAATA), which amplify a 150 bp fragment of the mitochondrial cytochrome b gene ^14^.

To optimize our amplification conditions, an initial PCR reaction was carried out on the control samples. We tested multiple annealing temperature ranging from 50 °C to 60 °C. Thermocycling conditions were set as follows: 94 °C for 5 minutes, followed by 35 cycles of each 94 °C for 30 seconds, 50 °C to 60 °C for 60 seconds (annealing step) and 72 °C for 60 seconds with a final extension at 60 °C for 45 minutes. To assess the sensitivity of these primers at low DNA quantities, dilutions were carried out on the Toronto control samples with concentrations of 0.5 ng/μl, 0.05 ng/μl, 0.005 ng/μl, and 0.0005 ng/μl. A 20 μl polymerase chain reaction (PCR) was prepared containing 2 μl of PCR Buffer (10X), 2 μl of 2 mM dNTPs, 1.33 μl of 3 μg/μl BSA, 0.8 μl of 50 mM MgCl2, 0.4 μl of each 10 μM primer, 0.2 μl of 5 U/μl Taq enzyme, 10.87 μl of ultra-pure DNase/RNase-free distilled water, and 2 μl of DNA extract. Along with the known snow leopard samples, a “no DNA template” negative control was included. PCR products were visualized under ultraviolet light on a 1.5% agarose gel stained with ethidium bromide. As all control samples amplified well, the optimal annealing temperature for the amplification of field samples was selected based on the brightness of the control samples that were at the lowest DNA concentrations.

DNA extracted from field samples was amplified using the same PCR and thermocycling conditions, at the identified optimal annealing temperature of 55 °C. Because the quantity of DNA and PCR inhibitors vary with each sample, a series of dilutions was prepared for each sample (1 in 5, 1 in 50, 1 in 500, 1 in 5000). The dilution for which amplification was most successful was selected for subsequent analyses. The hair controls were run at the quantities listed above and included for comparison on each gel. Forward and reverse sequences were obtained by using BigDye Terminator Cycle Sequencing Kit version 3.1 (Applied Biosystems, Inc.). Sequencing was performed on an automated DNA sequencer (ABI 3730; Applied Biosystems, Inc.). Sequences were analyzed using the phylogenetic software MEGA ^15^, and the NCBI BLAST database for species identification.

### Sex identification

Sex identification was performed using felid-specific primers (Zn-finger F: AAGTTTACACAACCACCTGG and R: CACAGAATTTACACTTGTGCA) targeting a zinc-finger region of the x and y-chromosomes ^16^. A fluorescent dye-label (HEX) was applied to the forward primer as the product size of Zfx at 166 base-pairs and 163 for Zfy were too close together to be distinguished by agarose gel ^16^. The 10 µL Zn-finger PCR reaction was prepared containing of 2 µL of 5X reaction buffer (Promega), 0.8 µL of 2.0 mM MgCl2, 1µL of 0.2 mM of each dNTP, 0.1 µL of each primer at 10 µM, 0.67 µL of 0.2 µg/µL BSA, 0.1µl of 5U/µl Taq polymerase(Promega) and 2µL of DNA. The PCR profile was 94 °C / 5 min, [94 °C / 1 min, 59 °C / 1 min, 72 °C / 1 min] x 34 cycles. PCR products were run in the ABI 3730 system. For each test, a “no DNA template” negative control was included. Positive control samples were included using high quality tissue from known cougar (*Puma concolor*) males and females, as well as scat and hair samples from known male and female snow leopards.

### Individual genotyping

Seven microsatellite loci (PUN100, PUN124, PUN132, PUN225, PUN229, PUN327 and PUN935 ^17^) were genotyped when DNA samples were identified as snow leopard DNA (details on primer sequences, labelling and annealing temperature are given in Table S2). A 10 μl PCR was prepared containing 2.0 μl of PCR Buffer (5X), 1.0 μl of 2 mM dNTPs, 1.0 μl of 3 μg/μl BSA, 0.6 μl of 25 mM MgCl2, 0.3 μl of each 10 μM primer, 0.1 μl of 5U/μl Taq enzyme, 2.7 μl of autoclaved ddH_2_O, and 2 μl of DNA extract. For each sample, PCR amplifications were run separately with similar conditions for each locus, except for PUN894. Thermocycling conditions set at 95 °C for 10 minutes, followed by 35 cycles at 95 °C for 15 seconds, 55 °C, and 72 °C for 60 seconds. An annealing temperature of 60 °C was used for PUN894. To verify amplification, PCR products were visualized under ultraviolet light on a 1.5% agarose gel stained with ethidium bromide. Samples for which no amplification was visible on the gel were amplified again with the total number of cycles increased to 45. The amplified products were then separated by size using capillary electrophoresis on an ABI 3730 DNA Analyzer system, and then sized using GeneMarker v1.9 software (SoftGenetics). We included hair and scat samples from one known individual with each ABI 3730 injection to ensure consistency in allele size and included positive and negative controls with each analysis. In order to prevent genotyping error, serial dilutions were conducted. This technique allows avoiding losing or miss-assigning alleles when signals from stock solutions are too strong. Under these circumstances, samples are unscorable because of wide peaks. When stock solutions that provided these wide peaks are diluted, peaks become clear, and the samples scorable. Serial dilutions also provide an opportunity to control for artifacts or errors in morphology from the ABI at different signal strengths. In this study, serial dilutions of samples that were overloaded on the ABI when their stock solution was used were set at 1/2, 1/5, 1/10, and 1/20 of the stock solution.

### Assessment of genotyping error

Genotyping error rate was estimated as the total number of allele call changes / total number of loci scored ^18^. All collected samples were analyzed at 7 microsatellite loci and at one sex locus (see sections Sex identification and Individual genotyping), for a total of 8 loci. Of all our samples, 61 were missing two or fewer of these eight loci, and were used in the subsequent GENECAP analysis ^19^ to identify specific individuals. Of these 61 samples, 51 were grouped into 7 individual profiles after close examination of allele peak scores for samples with similar genotypes. Because some samples had missing loci, the total number of loci scored for these 51 samples was 394 out of 408 possibilities (51 x 8 = 408). Upon close review of allele peaks scores by the technicians to identify potential allele drop-out or other quality issues such as inconsistent peak strength, morphology, and stutter, 7 changes were made to sex, one to PUN124, one to PUN132, 6 to PUN225, three to PUN235, and two to PUN299, for a total of 20 changes.

### Distribution of pairwise genetic distances according to kinship

Monitoring individuals requires the ability to discriminate them based on their respective genotypes. To do so, we first estimated the probability that two unrelated individuals would have the same genotype (*P*_*uni*_) using the following formula:

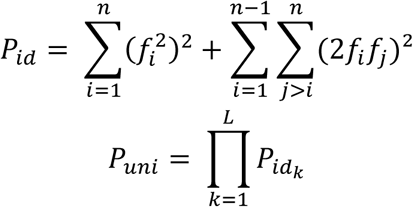

where P_*id*_ is the probability that two unrelated individuals have the same genotype at a given locus, *n* is the number of alleles at a given locus, *f*_*i*_ is the frequency of allele *i*, and *L* is the number of loci analyzed.

Although it is simple to compute the probability that two unrelated individuals have the same genotype, it is more difficult to compute the expected distribution of the genetic distance, that is the number of different alleles between two individuals, according to their kinship (*i.e.* unrelated, parent-descendant, full siblings, half siblings). For this study, an R script was written to compute these distributions using simulations (script_GenerateFamiliesIndividualRealFreqs-SupMaterials-Panthera.R in GitHub “https://github.com/jmorode/OSI-Panthera_genetics_scripts”). Pairs of genotypes were generated for different kinships. For two unrelated individuals, at each locus, the genotype of each individual was generated by randomly sampling two alleles, taking into account allele frequencies in the population. For a parent and its descendant, the genotype of the parent was generated as described above, and the genotype of the descendant was generated by sampling at random one allele from this parent and the other allele taking into account allele frequencies in the population. For full siblings, two unrelated parents were first generated; then, each descendant was generated by randomly sampling one allele from each parent. For half siblings, one mother and two unrelated fathers were first generated; then, one descendant was generated by randomly sampling one allele from the mother and one allele from one father. The other descendant was generated by randomly sampling one allele from the same mother and one allele from the other father. For each kind of relationship, one million simulations were performed to estimate the empirical distribution of pairwise distances based on kinship. All data analyses were carried out in the R statistical environment (version 3.4.3)^20^.

### Estimation of genealogical relationships

The software ML Relate ^21^ was used to find evidence of relatedness between snow leopards. The accuracy of inferred genealogical relationships by the software was evaluated using simulated families of individuals of known genotypes (script_GenerateIndividualRealFreqs-SupMaterials-Panthera.R in GitHub “https://github.com/jmorode/OSI-Panthera_genetics_scripts”). Two thousand families were generated using the above script. Each family was composed by a mother, two unrelated fathers, two full siblings and two pairs of half siblings. This family composition allowed testing the four relationships assessed by ML Relate: parent-offspring, full siblings, half siblings and unrelated. The genotypes of the members of these 2,000 families were written in an input file for ML Relate. For each pair of individuals in each family, the known relationship was compared with the one inferred by ML Relate. In each family, 6 unrelated individuals, 1 pair of full siblings, 2 pairs of half siblings and 6 pairs of parent-offspring were expected. The percentage of known relationships found by ML Relate over the 2,000 families was interpreted as an estimation of the accuracy of this software when identifying genealogical relationships in our observed population of snow leopards.

## Results

### Scat genotypes

DNA was extracted from 137 putative snow leopard scat samples. A snow leopard specific fragment of the *cytb* gene was amplified for 107 of these samples. The genotyping error rate, calculated from samples with two or fewer missing loci among 8 loci (7 microsatellites loci and 1 sex locus), was estimated at 20/394 = 5%.

Among those 107 samples, 51 samples were successfully genotyped at the 7 microsatellite loci. The other samples had missing data, 13 with one missing microsatellite locus, 5 with two missing microsatellite loci, and 38 with between 3 and 6 missing microsatellite loci. Samples genotyped with no more than one missing microsatellite locus are listed in Table 1 (the complete set of genotypes can be found in the file **supp_mat_data1_Genotypes.xls** in supplementary material). Twenty eight unique genotypes were identified, potentially corresponding to the same number of individual snow leopards. Seven of them were sampled several times (Table 1 and Fig. 2).

**Table 1.** Scat genotypes (no missing data or only at one locus)

| Individual | Sex | Locus |  |  |  |  |  |  |  |  |  |  |  |  |  | Number of recaptures |
| --- | --- | --- | --- | --- | --- | --- | --- | --- | --- | --- | --- | --- | --- | --- | --- | --- |
|  |  | PUN100 |  | PUN124 |  | PUN132 |  | PUN225 |  | PUN229 |  | PUN327 |  | PUN935 |  |  |
| 33_2014 | Female | 87 | 87 | 90 | 90 | 120 | 122 | 179 | 179 | 106 | 110 | 78 | 86 | 115 | 115 | 1 |
| 1_2012 | Female | 87 | 89 | 90 | 90 | 120 | 122 | 177 | 177 | 110 | 110 | 86 | 86 | 115 | 115 | 1 |
| SL6 / 4_2012 | Female | 87 | 89 | 90 | 90 | 120 | 122 | 177 | 179 | 110 | 110 | 86 | 86 | 115 | 115 | 8 |
| 26_2013 | - | 87 | 89 | 90 | 90 | 120 | 122 | 179 | 179 | 110 | 110 | 86 | 86 | 115 | 115 | 1 |
| 2_2015 | Male | 87 | 89 | 90 | 96 | 118 | 122 | 177 | 181 | 106 | 110 | 78 | 86 | 119 | 119 | 1 |
| 13_2015 | Male | 87 | 89 | 90 | 96 | 120 | 120 | 181 | 181 | 106 | 106 | 78 | 86 | 119 | 119 | 1 |
| 35_2013 | Male | 87 | 89 | 90 | 96 | 120 | 122 | 177 | 181 | 106 | 110 | 78 | 86 | 119 | 119 | 1 |
| 42_2013 | Female | 87 | 89 | 90 | 96 | 120 | 122 | 177 | 181 | 110 | 110 | 78 | 86 | 119 | 119 | 1 |
| 34_2013 | Male | 87 | 89 | 90 | 96 | 120 | 122 | 181 | 181 | 106 | 110 | 78 | 86 | 119 | 119 | 1 |
| 17_2012 | Male | 89 | 89 | 90 | 90 | 120 | 122 | 179 | 179 | ** | ** | 86 | 86 | 115 | 115 | 1 |
| 6_2015 | Female | 89 | 89 | 90 | 96 | 120 | 122 | 175 | 181 | 110 | 110 | 78 | 86 | 119 | 119 | 1 |
| 2_2013 | Male | 89 | 89 | 90 | 96 | 120 | 122 | 177 | 177 | 104 | 108 | 78 | 88 | 115 | 119 | 1 |
| SL1 / 12_2012 | Male | 89 | 89 | 90 | 96 | 120 | 122 | 177 | 177 | 106 | 106 | 78 | 88 | 115 | 119 | 15 |
| 23_2012 | Male | 89 | 89 | 90 | 96 | 120 | 122 | 177 | 181 | 106 | 106 | 78 | 88 | 115 | 119 | 1 |
| SL2 / 2_2014 | Male | 89 | 89 | 90 | 96 | 120 | 122 | 177 | 181 | 106 | 110 | 78 | 86 | 119 | 119 | 4 |
| 14_2011 | - | 89 | 89 | 90 | 96 | 120 | 122 | ** | ** | 106 | 106 | 78 | 88 | 115 | 119 | 1 |
| SL3 / 6_2014 | Male | 89 | 89 | 100 | 102 | 112 | 120 | 177 | 179 | 106 | 110 | 78 | 86 | 115 | 119 | 3 |
| 3_2015 | Male | 89 | 91 | 90 | 96 | 120 | 122 | 177 | 181 | 106 | 110 | 78 | 86 | 119 | 119 | 1 |
| SL5 / 13_2011 | Female | 89 | 91 | 90 | 100 | 112 | 118 | 175 | 175 | 110 | 110 | 78 | 86 | 115 | 119 | 3 |
| 5_2015 | Female | 89 | 91 | 90 | 100 | 112 | 122 | 175 | 179 | 106 | 108 | 78 | 86 | 119 | 119 | 1 |
| SL4 / 17_2013 | Male | 89 | 93 | 90 | 90 | 120 | 122 | 175 | 179 | 106 | 110 | 86 | 86 | 119 | 119 | 11 |
| 31_2014 | Female | 89 | 93 | 90 | 96 | 122 | 122 | 175 | 179 | ** | ** | 86 | 86 | 119 | 119 | 1 |
| 11_2012 | Female | 89 | 93 | 90 | 100 | 112 | 122 | 175 | 177 | ** | ** | 78 | 86 | 119 | 119 | 1 |
| 20_2014 | Male | 89 | 93 | ** | ** | 120 | 122 | 175 | 179 | 106 | 110 | 86 | 86 | 119 | 119 | 1 |
| 8_2015 | - | 91 | 95 | 90 | 90 | 120 | 122 | 177 | 177 | 108 | 108 | 86 | 86 | 119 | 119 | 1 |
| 7_2011 | - | 93 | 93 | 90 | 90 | 120 | 120 | 181 | 181 | ** | ** | 86 | 86 | 115 | 119 | 1 |
| SL7 / 6_2011 | - | 93 | 93 | 90 | 90 | 120 | 122 | 175 | 181 | 110 | 110 | 78 | 86 | 115 | 119 | 3 |
| 22_2013 | - | 93 | 93 | 90 | 90 | 122 | 122 | 173 | 175 | 106 | 106 | 86 | 86 | ** | ** | 1 |

If they were not recaptured, individual names are their sampling number. If they were recaptured, they have a name SLx / sampling number of the oldest sample recovered.

**Figure 2.**
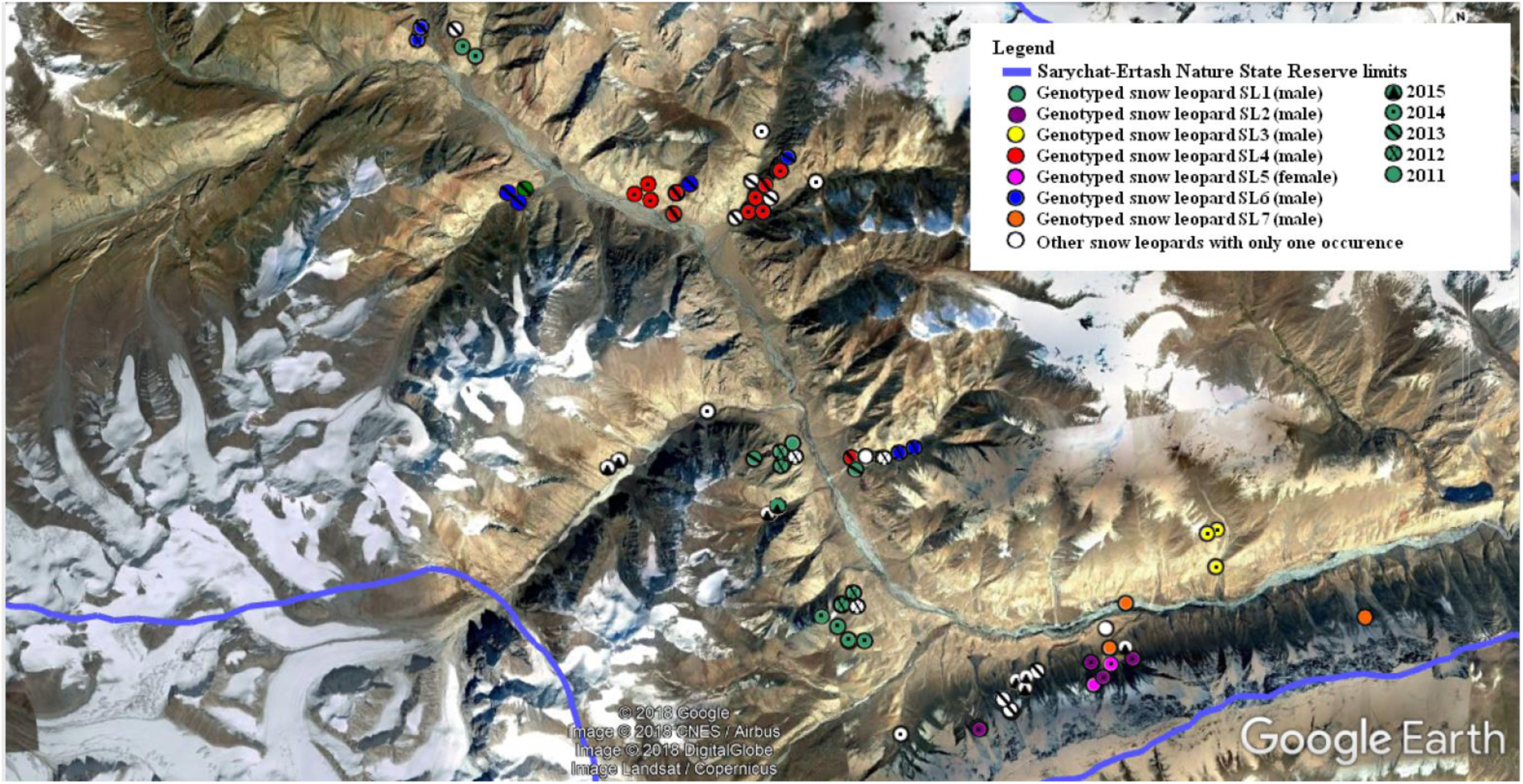
Location of the genotyped snow leopard scats in the Sarychat-Ertash State Reserve from 2011 to 2015.

### Distributions of pairwise genetic distances according to genealogical relationships

Allele frequencies and expected heterozygosity were estimated using all snow leopard samples (Table 2). Pairwise distances between the different genotypes were also computed (Table 3 and Fig. 3). Most pairwise distances were between five and ten differences. However, some pairs of genotypes differed by only one or two alleles (9 pairs and 17 pairs, respectively).

**Table 2.** Allele frequencies and expected heterozygosity

|  | **Locus** |  |  |  |  |  |  |  |  |  |  |  |  |  |
| --- | --- | --- | --- | --- | --- | --- | --- | --- | --- | --- | --- | --- | --- | --- |
|  | PUN100 |  | PUN124 |  | PUN132 |  | PUN225 |  | PUN229 |  | PUN327 |  | PUN935 |  |
| **Number of alleles** | 5 |  | 6 |  | 7 |  | 8 |  | 4 |  | 4 |  | 3 |  |
| **Allele size and frequency** | 87 | 0,179 | 90 | 0,661 | 110 | 0,017 | 171 | 0,016 | 104 | 0,021 | 78 | 0,333 | 115 | 0,339 |
|  | 89 | 0,554 | 96 | 0,210 | 112 | 0,083 | 173 | 0,016 | 106 | 0,396 | 86 | 0,600 | 119 | 0,625 |
|  | 91 | 0,089 | 100 | 0,081 | 114 | 0,017 | 175 | 0,210 | 108 | 0,083 | 88 | 0,067 | 127 | 0,036 |
|  | 93 | 0,161 | 102 | 0,016 | 116 | 0,017 | 177 | 0,323 | 110 | 0,500 |  |  |  |  |
|  | 95 | 0,018 | 106 | 0,016 | 118 | 0,050 | 179 | 0,210 |  |  |  |  |  |  |
|  |  |  | 108 | 0,016 | 120 | 0,400 | 181 | 0,226 |  |  |  |  |  |  |
|  |  |  |  |  | 122 | 0,417 |  |  |  |  |  |  |  |  |
| **He** | 0,63 |  | 0,51 |  | 0,66 |  | 0,76 |  | 0,59 |  | 0,52 |  | 0,49 |  |

**Table 3.**
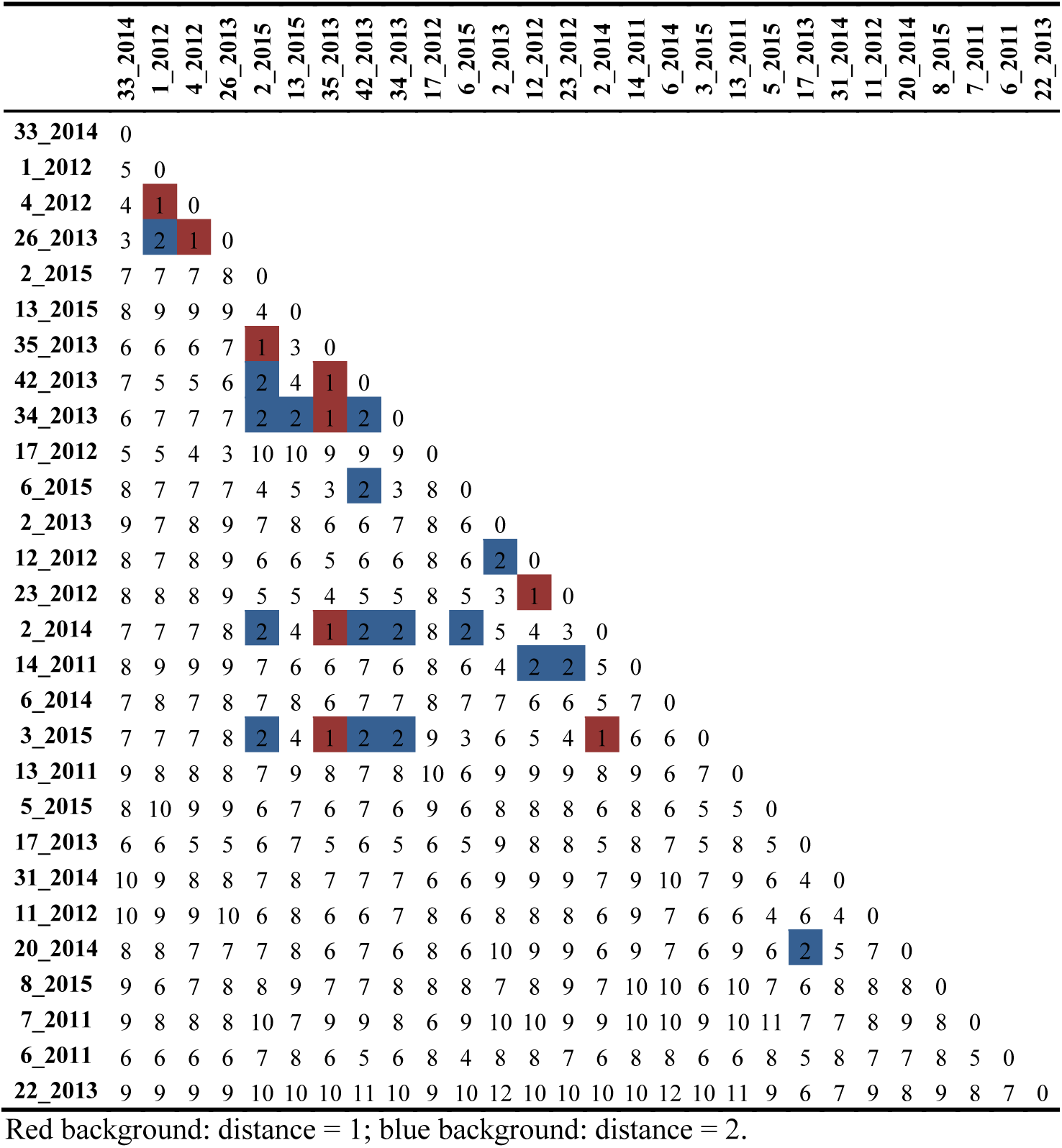
Pairwise distances between genotypes

**Figure 3.**
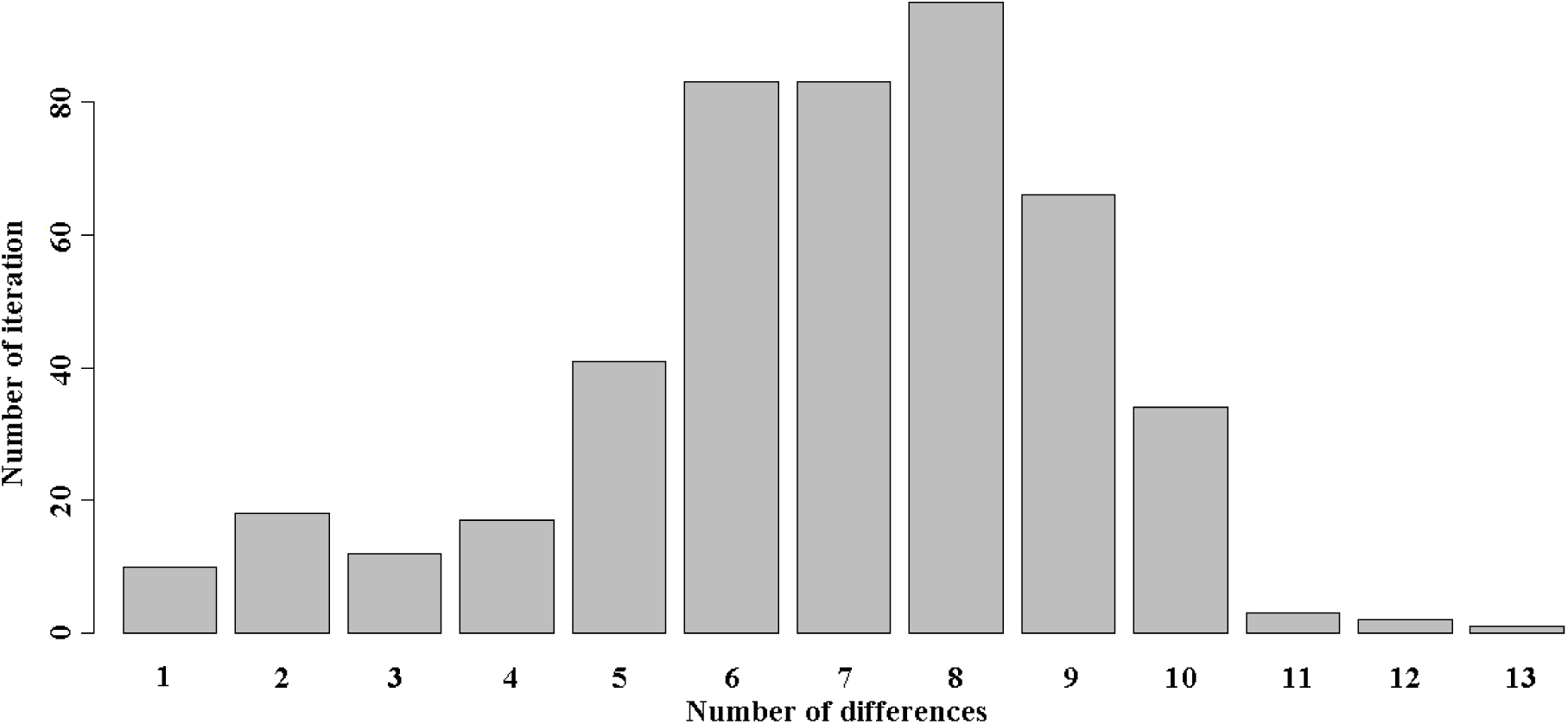
Pairwise distances between genotypes.

Considering the estimated allele frequencies in the population, the probability that two unrelated individuals could have the same genotype (P_*uni*_) was very low (2.73 × 10^−5^). When considering only pairwise distances with a probability above 5%, our simulations showed that pairwise distances between two individuals would be in the ranges of: i) 2 to 6 differences for parent offspring pairs; ii) 1 to 6 differences for full siblings; iii) 3 to 8 differences for half siblings; and iv) 4 to 9 differences for unrelated individuals (Fig. 4).

**Figure 4.**
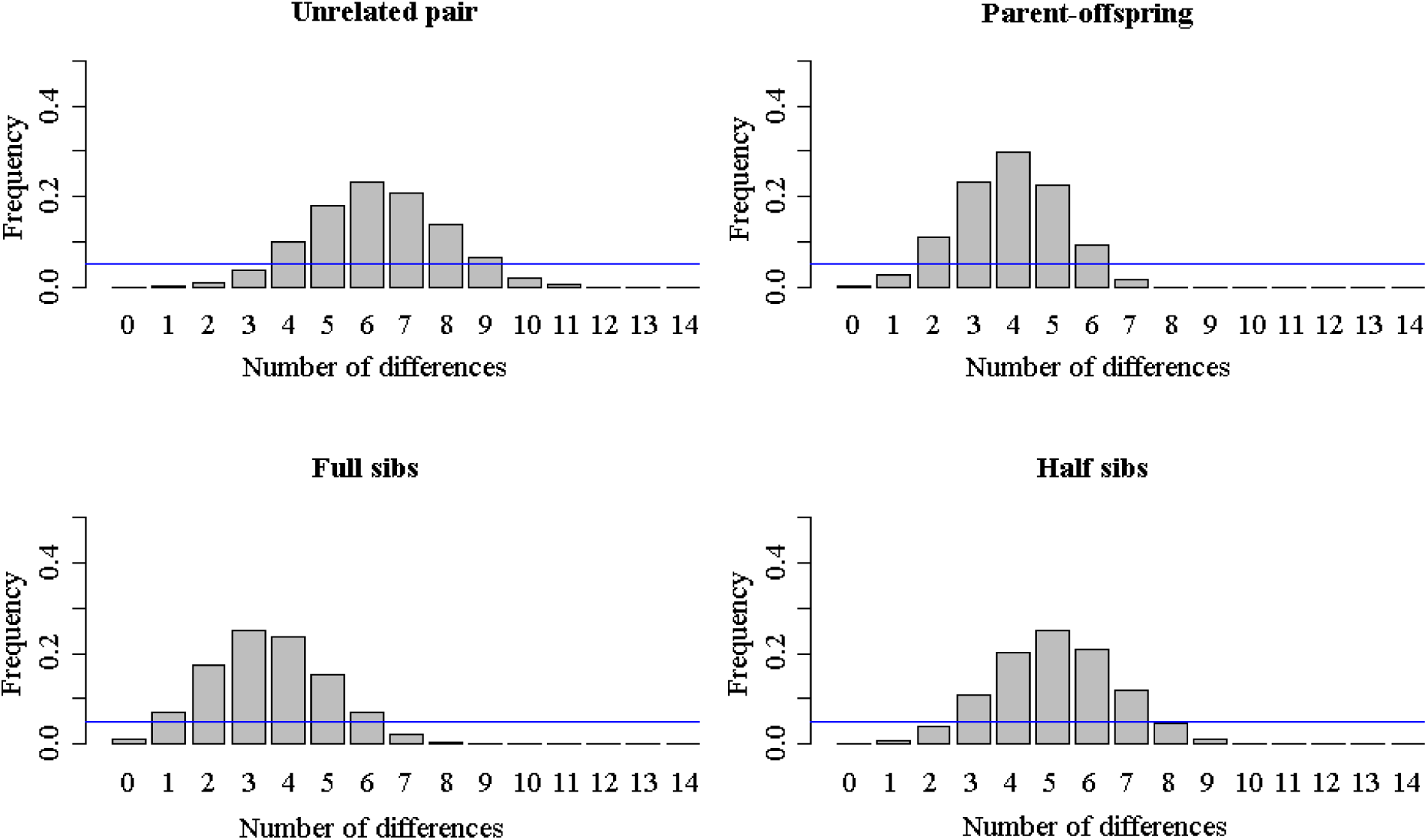
Expected distribution of pairwise genetic distance according to genealogical relationship. Genetic distance is defined as the number of allele differences between two individuals.

### Minimum number of snow leopards

Although collected samples were carefully genotyped, genotyping errors are possible when the DNA is degraded, which is often the case with scat samples (see “Scat genotypes” above). Assuming different unknown amount of errors due to allelic dropout and null alleles, we estimated the number of snow leopards genotyped. Under the most optimistic hypothesis that assumed no genotyping errors, and taking into account the samples genotyped at all microsatellite loci and the samples with one missing microsatellite locus, 28 snow leopards were identified. Under a much more pessimistic hypothesis that assumed that up to two differences are systematically the result of genotyping errors and taking into account only samples for which a complete genotype was available, the number of snow leopards genotyped dropped to 11 (Fig. 5). In any case, the number of identified individuals grew steadily from 2011 to 2015, with no evidence of saturation (Fig. 5).

**Figure 5.**
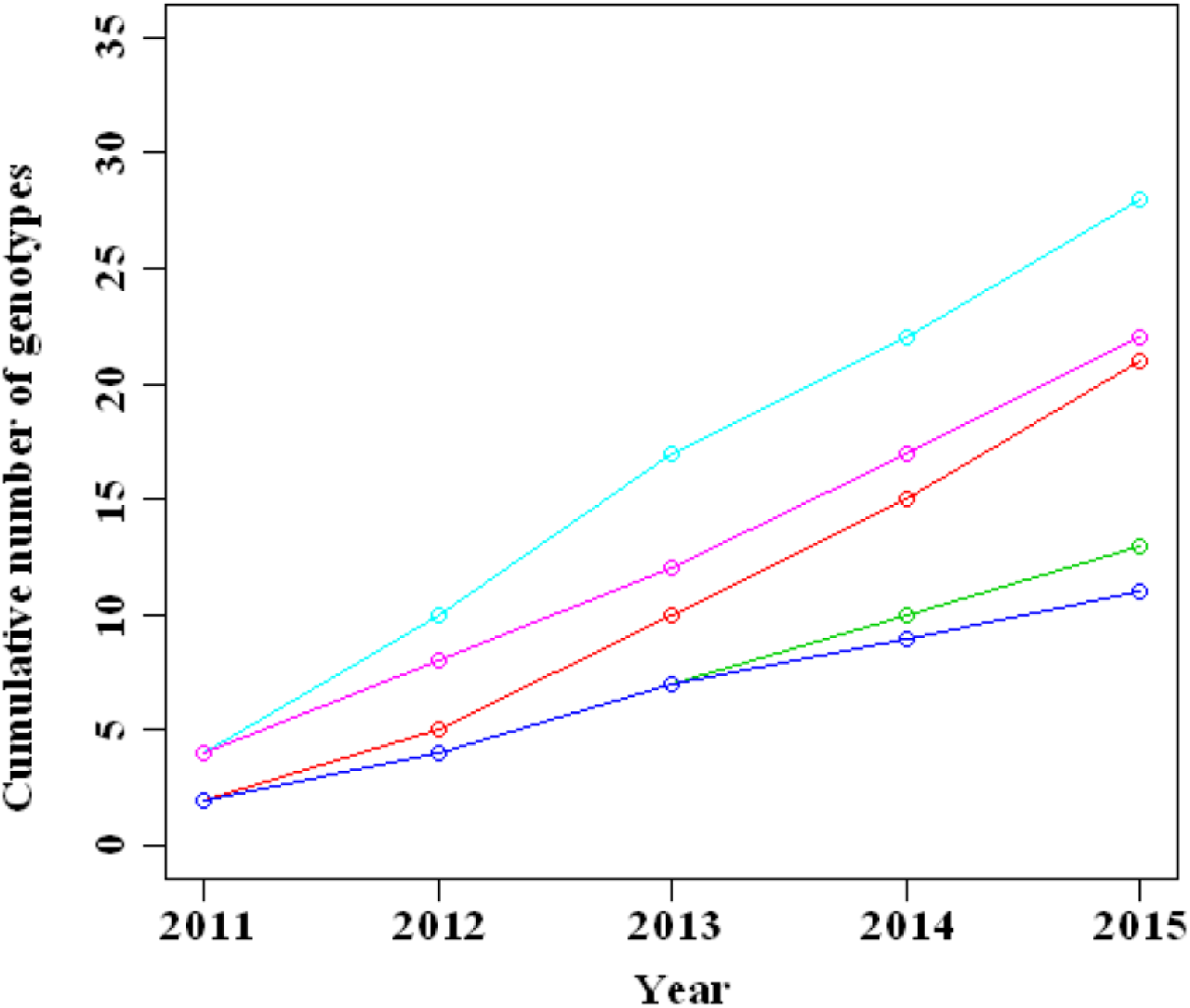
Cumulative number of snow leopard different genotypes identified from 2011 to 2015. Red: complete genotypes only; green: complete genotypes with the possibility of one genotyping error; blue: complete genotypes with the possibility of two genotyping errors; light blue: complete genotypes and genotypes with one locus not genotyped; purple: complete genotypes and genotypes with one locus not genotyped and with the possibility of one genotyping error; yellow: complete genotypes and genotypes with one locus not genotyped and with the possibility of two genotyping errors.

### Relationships between snow leopards

Relationships between individual snow leopards identified through genotyping were estimated using ML Relate. With the 22 individuals that had complete genotypes (no missing microsatellite data), 16 parent-offspring, 31 full siblings and 13 half siblings relationships were found. When the genotypes with missing data at one microsatellite locus were added (29 individuals identified with one missing locus), 34 parent-offspring, 42 full siblings and 25 half siblings relationships were found. In both cases, each individual was related to between two and 12 other individuals (**Fig. S1** in supplementary material). Relationship matrices can be found in the file **supp_mat_data2_MLRelateOutput.xls** in supplementary material.

### Reliability of ML Relate

The assessment of ML Relate’s performance using simulated families showed that the program had a 62% accuracy, on average. Parent-offspring relationships were correctly found in 71% of cases, unrelated individuals in 69% of cases, full siblings in 48% of cases, and half siblings in only 20% of cases. When the relationship provided by ML Relate was incorrect (38% of the time), we found that: i) 32% of pairs of individuals indicated as unrelated by ML Relate included in fact 13% parent-offspring, 6% full siblings, and 13% half siblings; ii) 31% of pairs of individuals indicated as parent-offspring by ML Relate included in fact 3% of unrelated pairs, 22% of full siblings, and 6% of half siblings; iii) 10% of pairs of individuals indicated as full siblings by ML Relate included in fact 2% of unrelated pairs, 6% of parent-offspring, and 2% of half siblings; iv) 27% of pairs of individuals indicated as half siblings by ML Relate included in fact 11% of unrelated pairs, 11% parent-offspring, and 5% full siblings.

### Diachronic monitoring of individuals

Monitoring of individuals inside the reserve was performed using only scat samples for which six to seven microsatellite loci were genotyped. Assuming no genotyping errors, seven snow leopards (SL1 to SL7) were sampled more than once (Fig. 2). A total of 15 scats belonging to SL1 were collected between 2011 and 2015 in Bir Baital (P), Uch Baital (N), Chomoi (G), Saryetchki (Q), Eki Baital (O) and Orto-Bordu (E). This snow leopard was present across the main valley, and was able to cross the Ertash River, as its feces were collected from both river banks. A total of 4 scats belonging to SL2 were collected in 2014 in the southeastern part of SESR on the southern side of the Ertash River, in the Solomo (U) and Sirdibai (T) areas. A total of 3 scats belonging to SL3 were collected in 2014 in Jili Boulak (V). A total of 11 scats belonging to SL4 were collected in 2013 and 2014 in the core part of the main valley on the eastern side of the Ertash River (Kirk-choro (H), Gueuleu (L) and Sary etchki (Q)). A total of 3 scats belonging to SL5 were collected in 2011, 2014 and 2015 in Solomo (U). A total of 8 scats belonging to SL6 were collected in 2012 and 2013 in Bordu (C), Chomoi (G), Kirk-choro (H) and Gueuleu (L). Finally, a total of 3 scats belonging to SL7 were collected in 2011 in Solomo (U) and Orto Choi (X). The maps corresponding to each individual can be found in the **supplementary materials (maps SL1 to SL7)**.

## Discussion

Thanks to repeated OSI-Panthera expeditions, individual snow leopards were identified by genotyping scat samples collected over multiple years. Through this non-invasive capture-recapture method, it was possible to follow individuals’ movements. The mere existence of this study shows that citizen science expeditions are a powerful way to gather field data over long periods of time and across large areas when logistical and financial demands cannot be met via academic research grants ^22^.

Scat samples were often degraded when collected because of harsh weather and high altitude. Thus, we expected missing genotypes and allelic dropout ^23^. This hindered the optimization of the genotyping process, as DNA was sometimes not of good enough quality for individual identification. Nonetheless, with seven microsatellite loci, a minimum of 11 snow leopards were identified, seven of which were sampled multiple times and at different locations within the SESR.

New genotypes were steadily identified each year with no evidence of saturation, which could be explained by one or several of the following: *i*) not enough scats were collected to identify most snow leopards in the sampled area; *ii*) the birth of cubs during the sampling period, which could have induced the sampling of new individuals; *iii*) adults and subadults coming regularly from outside the sampling area into SESR. A more extensive survey is necessary to disentangle the importance of these factors.

The most conservative estimation of the number of snow leopards identified in this study, 11 individuals, is close to the number of 15 individuals identified via camera trapping in 2014 ^4^, and to the 18 individuals identified with genetic markers across a 1,341 km² area in 2009^4^.

When including complete genotypes and genotypes with only one missing microsatellite locus, 14 males and 9 females were identified (Table 1). Although this sex ratio (1.55:1) is not significantly different from a 1:1 sex ratio (chi-squared test, p = 0.33), it will be important to assess its evolution over a longer period of time in the SESR, as snow leopard sex ratio can be highly dynamic, which could affect the renewal potential of the population ^24^.

### Genealogical relationships

Many relationships between individuals were found by ML Relate (60relationships when complete genotypes were used, and 101 when genotypes containing up to one missing locus were used). All individual snow leopards were related to at least two other individuals. Some individuals were shown to have ten relationships, although some of them were in fact inconsistent (for example, individual 2_2015 was identified by ML Relate as being both full sibling with 13_2015 and 35_2013, which were indicated as being parent-offspring). Simulated pairs of individuals with known relatedness showed that the overall accuracy of ML Relate was low (62%), indicating that the number of loci used in this study was not large enough to reliably identify the relationships between snow leopards. This is particularly the case for estimated full siblings and half sibling relationships, which showed an accuracy level of only 48% and 20%, respectively. In contrast, parent offspring and unrelated individuals were identified more accurately (71% and 69%, respectively). A better knowledge of the relationships between individuals could be valuable information to estimate the level of inbreeding and the number of descendants per male and female, but more data are necessary to achieve this goal.

Although there was a lack of accuracy when identifying genealogical relationships with ML Relate, the expected distribution of pairwise distances between individuals suggests that if only one or two differences were found between two genotypes, it would be likely that these individuals are related. One or two allelic differences between two individuals would most likely represent a close relationship, or the genotype of only one individual with genotyping errors. Ten pairs of genotypes showed only one allelic difference, and 18 showed two allelic differences. Among these pairs, two pairs with one difference and 5 with two differences involved both a male and a female, indicating that these could not have been the result of genotyping the same individual with errors. In addition, all these pairs of genotypes were identified as full siblings by ML Relate. In contrast, there was an excess of same sex pairs among the pairs of genotypes with one or two allelic differences (8/10 and 14/18 respectively). Under a 1:1 sex ratio, we would expect that half the full sibling pairs are of the same sex. The excess of same sex pairs may thus indicate that some closely related genotypes are the result of genotyping errors. This suggests that at least the 7 male-female pairs observed out of the 28 closely related pairs of genotypes represent closely related snow leopards, which may be full siblings, as inferred by ML Relate.

### Diachronic monitoring of individuals

Some individuals were recaptured several times, across multiple years. Two individuals, SL1 (male) and SL6 (male), were found throughout the main valley of the SESR core, across a large territory. Male and female territories overlapped in the study area, as indicated in previous studies ^4^. SL7 was sampled three times in 2011, at three very close locations, and was never sampled again; expeditions may have not cross its path again, his main territory could have moved outside the reserve, or he could have died.

### Conclusion

In conclusion, our analyses allowed a better understanding of the snow leopard population in the SESR from 2011 to 2015. In the near future, we could refine our analyses by continuing scat sampling efforts, ground-truthing relationships by sampling scats around known den sites, and genotyping at more loci. Using additional approaches such as camera trapping to identify individuals and scat location would also help refine snow leopard identification. These improvements to the current methodology would provide a deeper understanding of the SESR snow leopard population dynamics, and thus, offer further inputs for the conservation of this species in Kyrgyzstan.

## Acknowledgements

We would like to warmly thank the support of every volunteer of the OSI-Panthera expeditions along with all the scientific educators and rangers of the reserve who made this study possible. We would also like to thank Rob Moyes and Eric “Kuba” Ash at Jungle Cat World, the Toronto zoo Animal Care staff for securing control samples, and Emily Walker and Tasnova Kahn for their help with the processing of our first field samples. Special thanks to Muktar Musaev (Sarychat-Ertash Reserve director), Jean-Marc Elalouf and Jose Utge (MNHN), for their help in improving our sampling and genotyping protocols. This study was supported by the Natural Resource DNA and Profiling Forensic Center (19,919.33 $CA), the Endangered Species Reserve Fund Program of the Toronto Zoo (1,000 $CA), the PPG Conservation and Sustainability Fund of the Pittsburgh Zoo & PPG Aquarium (1,500 $US), and by a Faculty Research grant from the Canadian Mennonite University (2,500 $CA). This study was funded thanks to the citizens which took part in the many expeditions.

## Author contributions

J.R., A.P., S.R., A.L.C., A.O., B.C. designed the study. J.R, A.P., S.R., A.L.C., A.O., B.C. and A.V. collected the samples. A.P., B.W., M.H. and N.T.X. performed PCR amplifications, sequencing and scoring of the microsatellite markers. J.R., J.F. and D.C. analyzed the data. J.R., A.P. and D.C. wrote the manuscript. All authors revised the manuscript and approved the final version.

## Ethics approval and consent to participate

Scat and hair samples from captive animals were collected non-invasively on the floor with the permission of the Animal Care director of the Toronto Zoo.

An authorization to collect feces was given by the head of Sarychat Ertash State Reserve. In the OSI participation conditions (http://www.vacances-scientifiques.com/Conditions-de-Participation.html), it is stated that data gathered by participants during these expeditions will be used for scientific purposes.

## Data Availability

Data are available in Supplementary Data S1.xlsx and Supplementary Data S2.xlsx

