## Supplemental data 2.1 - MLRelate Output for "Diachronic monitoring of snow leopards at Sarychat-Ertash State Reserve (Kyrgyzstan) through scat genotyping: a pilot study" (from another version of this preprint: version 1, 2019-11-16)

**Full genotypes (no missing data)**

| | 33_2014 | 1_2012 | SL6 / 4_2012 | 26_2013 | 2_2015 | 13_2015 | 35-2013 | 42_2013 | 34_2013 | 6_2015 | 2_2013 | SL1 / 12_2012 | 23_2012 | SL2 / 2_2014 | SL3 / 6_2014 | 3_2015 | SL5 / 13_2011 | 5_2015 | SL4 / 17_2013 | 23_2013 | 8_2015 | SL7 / 6_2011 | 10_2011 |
|---|---|---|---|---|---|---|---|---|---|---|---|---|---|---|---|---|---|---|---|---|---|---|---|
| **33_2014** | - | | | | | | | | | | | | | | | | | | | | | | |
| **1_2012** | PO | - | | | | | | | | | | | | | | | | | | | | | |
| **L6 / 4_201** | PO | FS | - | | | | | | | | | | | | | | | | | | | | |
| **26_2013** | FS | FS | FS | - | | | | | | | | | | | | | | | | | | | |
| **2_2015** | U | U | U | U | - | | | | | | | | | | | | | | | | | | |
| **13_2015** | U | U | U | U | FS | - | | | | | | | | | | | | | | | | | |
| **35-2013** | U | U | U | U | FS | PO | - | | | | | | | | | | | | | | | | |
| **42_2013** | U | HS | U | U | FS | FS | FS | - | | | | | | | | | | | | | | | |
| **34_2013** | U | U | U | U | FS | FS | FS | FS | - | | | | | | | | | | | | | | |
| **6_2015** | U | U | U | U | HS | HS | PO | FS | PO | - | | | | | | | | | | | | | |
| **2_2013** | U | U | U | U | U | U | U | U | U | U | - | | | | | | | | | | | | |
| **L1 / 12_20** | U | U | U | U | U | U | HS | U | U | U | FS | - | | | | | | | | | | | |
| **23_2012** | U | U | U | U | U | PO | PO | U | PO | U | FS | FS | - | | | | | | | | | | |
| **L2 / 2_201** | U | U | U | U | FS | PO | FS | FS | FS | FS | U | PO | FS | - | | | | | | | | | |
| **L3 / 6_201** | U | U | U | U | U | U | U | U | U | U | U | U | U | U | - | | | | | | | | |
| **3_2015** | U | U | U | U | FS | PO | FS | FS | FS | PO | U | HS | PO | FS | U | - | | | | | | | |
| **L5 / 13_20** | U | U | U | U | U | U | U | U | U | U | U | U | U | U | HS | U | - | | | | | | |
| **5_2015** | U | U | U | U | U | U | U | U | U | U | U | U | U | U | PO | U | FS | - | | | | | |
| **L4 / 17_20** | U | U | U | HS | U | U | U | U | U | PO | U | U | U | U | U | U | U | HS | - | | | | |
| **23_2013** | U | U | U | U | U | U | U | U | U | HS | U | U | U | U | U | U | U | HS | FS | - | | | |
| **8_2015** | U | U | U | U | U | U | U | U | U | U | U | U | U | U | U | HS | U | HS | U | U | - | | |
| **L7 / 6_201** | U | U | U | U | U | U | U | U | U | FS | U | U | U | U | U | U | U | U | PO | HS | U | - | |
| **10_2011** | U | U | U | U | U | U | U | U | U | U | U | U | U | U | U | U | U | U | HS | U | U | FS | - |

| | Number |
|---|---|
| PO | 16 |
| FS | 32 |
| HS | 14 |

**Null alleles (output of MLRELATE) :**
Hardy-Weinberg test for heterozygote deficiency

This randomization test uses Monte Carlo randomization as described by Guo & Thompson (1992) and the 'U' test statistic described by Rousset and Raymond (1995).

NRandomizations = 1000000

| Locus | PValue |
|---|---|
| PUN100 | 0,1117 |
| PUN124 | 0,9323 |
| PUN132 | 0,9999 |
| PUN225 | 0,0289 |
| PUN229 | 0,0182 |
| PUN327 | 0,9873 |
| PUN935 | 0,0192 |

Interpretation: If the p value listed above is low, you can conclude that there is a heterozygote deficiency. Please note that null alleles can cause heterozygote deficiencies, but there are other possible explanations.
