## Supplemental data 2.2 - MLRelateOutput for "Diachronic monitoring of snow leopards at Sarychat-Ertash State Reserve (Kyrgyzstan) through scat genotyping: a pilot study" (from another version of this preprint: version 1, 2019-11-16)

**Full genotypes (no missing data) and genotypes with one locus with missing data**

| | 33_2014 | 1_2012 | SL6 / 4_2012 | 26_2013 | 2_2015 | 13_2015 | 35_2013 | 42_2013 | 34_2013 | 9_2015 | 17_2012 | 6_2015 | 2_2013 | SL1 / 12_2012 | 23_2012 | SL2 / 2_2014 | 14_2011 | SL3 / 6_2014 | 3_2015 | SL5 / 13_2011 | 5_2015 | SL4 / 17_2013 | 23_2013 | 31_2014 | 11_2012 | 20_2014 | 8_2015 | 7_2011 | SL7 / 6_2011 | 10_2011 | 22_2013 |
|---|---|---|---|---|---|---|---|---|---|---|---|---|---|---|---|---|---|---|---|---|---|---|---|---|---|---|---|---|---|---|---|
| 33_2014 | - | | | | | | | | | | | | | | | | | | | | | | | | | | | | | | |
| 1_2012 | PO | - | | | | | | | | | | | | | | | | | | | | | | | | | | | | | |
| SL6 / 4_2012 | PO | FS | - | | | | | | | | | | | | | | | | | | | | | | | | | | | | |
| 26_2013 | PO | FS | FS | - | | | | | | | | | | | | | | | | | | | | | | | | | | | |
| 2_2015 | U | U | U | U | - | | | | | | | | | | | | | | | | | | | | | | | | | | |
| 13_2015 | HS | U | U | U | FS | - | | | | | | | | | | | | | | | | | | | | | | | | | |
| 35_2013 | U | HS | U | U | FS | PO | - | | | | | | | | | | | | | | | | | | | | | | | | |
| 42_2013 | U | HS | HS | U | FS | FS | FS | - | | | | | | | | | | | | | | | | | | | | | | | |
| 34_2013 | HS | U | U | U | FS | FS | FS | FS | - | | | | | | | | | | | | | | | | | | | | | | |
| 9_2015 | U | U | U | U | U | U | U | U | U | - | | | | | | | | | | | | | | | | | | | | | |
| 17_2012 | FS | FS | PO | FS | U | U | U | U | U | U | - | | | | | | | | | | | | | | | | | | | | |
| 6_2015 | U | U | U | U | PO | HS | PO | FS | PO | U | U | - | | | | | | | | | | | | | | | | | | | |
| 2_2013 | U | U | U | U | U | U | U | U | U | FS | U | U | - | | | | | | | | | | | | | | | | | | |
| SL1 / 12_2012 | U | U | U | U | U | HS | PO | U | U | FS | U | U | FS | - | | | | | | | | | | | | | | | | | |
| 23_2012 | U | U | U | U | HS | PO | FS | U | PO | FS | U | U | FS | FS | - | | | | | | | | | | | | | | | | |
| SL2 / 2_2014 | U | U | U | U | FS | PO | FS | FS | FS | PO | U | FS | U | PO | FS | - | | | | | | | | | | | | | | | |
| 14_2011 | U | U | U | U | U | PO | U | U | U | FS | U | U | FS | FS | FS | PO | - | | | | | | | | | | | | | | |
| SL3 / 6_2014 | U | U | U | U | U | U | U | U | U | U | U | U | U | U | U | U | U | - | | | | | | | | | | | | | |
| 3_2015 | U | U | U | U | FS | PO | FS | FS | FS | U | U | PO | U | PO | FS | FS | U | U | - | | | | | | | | | | | | |
| SL5 / 13_2011 | U | U | U | U | HS | U | U | U | U | U | U | U | U | U | U | U | U | HS | U | - | | | | | | | | | | | |
| 5_2015 | U | U | U | U | U | U | U | U | U | U | U | U | U | U | U | U | U | PO | U | FS | - | | | | | | | | | | |
| SL4 / 17_2013 | U | U | HS | HS | U | U | U | U | U | U | HS | PO | U | U | U | U | U | U | U | U | U | - | | | | | | | | | |
| 23_2013 | U | U | U | U | U | U | U | U | U | U | HS | HS | U | U | U | U | U | U | U | U | U | FS | - | | | | | | | | |
| 31_2014 | U | U | U | U | U | U | U | U | U | U | HS | PO | U | U | U | U | U | U | U | U | HS | FS | FS | - | | | | | | | |
| 11_2012 | U | U | U | U | U | U | U | U | U | U | U | U | U | U | U | U | U | PO | U | PO | FS | HS | U | HS | - | | | | | | |
| 20_2014 | U | U | U | U | U | U | U | U | U | U | HS | PO | U | U | U | U | U | U | U | U | U | FS | FS | FS | PO | - | | | | | |
| 8_2015 | U | U | U | U | U | U | U | U | U | U | U | U | HS | U | U | U | U | U | HS | U | HS | U | U | U | U | U | - | | | | |
| 7_2011 | U | U | U | U | U | HS | U | U | U | U | U | U | U | U | U | U | U | U | U | U | U | HS | U | U | U | HS | U | - | | | |
| SL7 / 6_2011 | U | U | U | U | U | U | U | U | U | U | U | FS | U | U | U | U | U | U | U | U | U | PO | U | PO | HS | PO | U | PO | - | | |
| 10_2011 | U | U | U | U | U | U | U | U | U | U | U | U | U | U | U | U | U | U | U | U | U | U | U | U | U | U | U | FS | FS | - | |
| 22_2013 | U | U | U | U | U | U | U | U | U | U | U | U | U | U | U | U | U | U | U | U | U | PO | PO | PO | PO | PO | U | FS | PO | HS | - |

| | Number |
|---|---|
| PO | 36 |
| FS | 51 |
| HS | 28 |

**Null alleles (output of MLRELATE) :**
Hardy-Weinberg test for heterozygote deficiency

This randomization test uses Monte Carlo randomization as described by Guo & Thompson (1992)
and the 'U' test statistic described by Rousset and Raymond (1995).

NRandomizations = 1000000

| Locus | PValue |
|---|---|
| PUN100 | 0,121 |
| PUN124 | 0,941 |
| PUN132 | 1 |
| PUN225 | 0,013 |
| PUN229 | 0,001 |
| PUN327 | 0,966 |
| PUN935 | 0,057 |

Interpretation: If the p value listed above is low, you can conclude
that there is a heterozygote deficiency. Please note that null alleles
can cause heterozygote deficiencies, but there are other possible explanations.
