## Supplementary figures and images for "Diachronic monitoring of snow leopards at Sarychat-Ertash State Reserve (Kyrgyzstan) through scat genotyping: a pilot study" (from another version of this preprint: version 1, 2019-11-16)

### Supplemental Figure 1 _ relationships from MLRelate

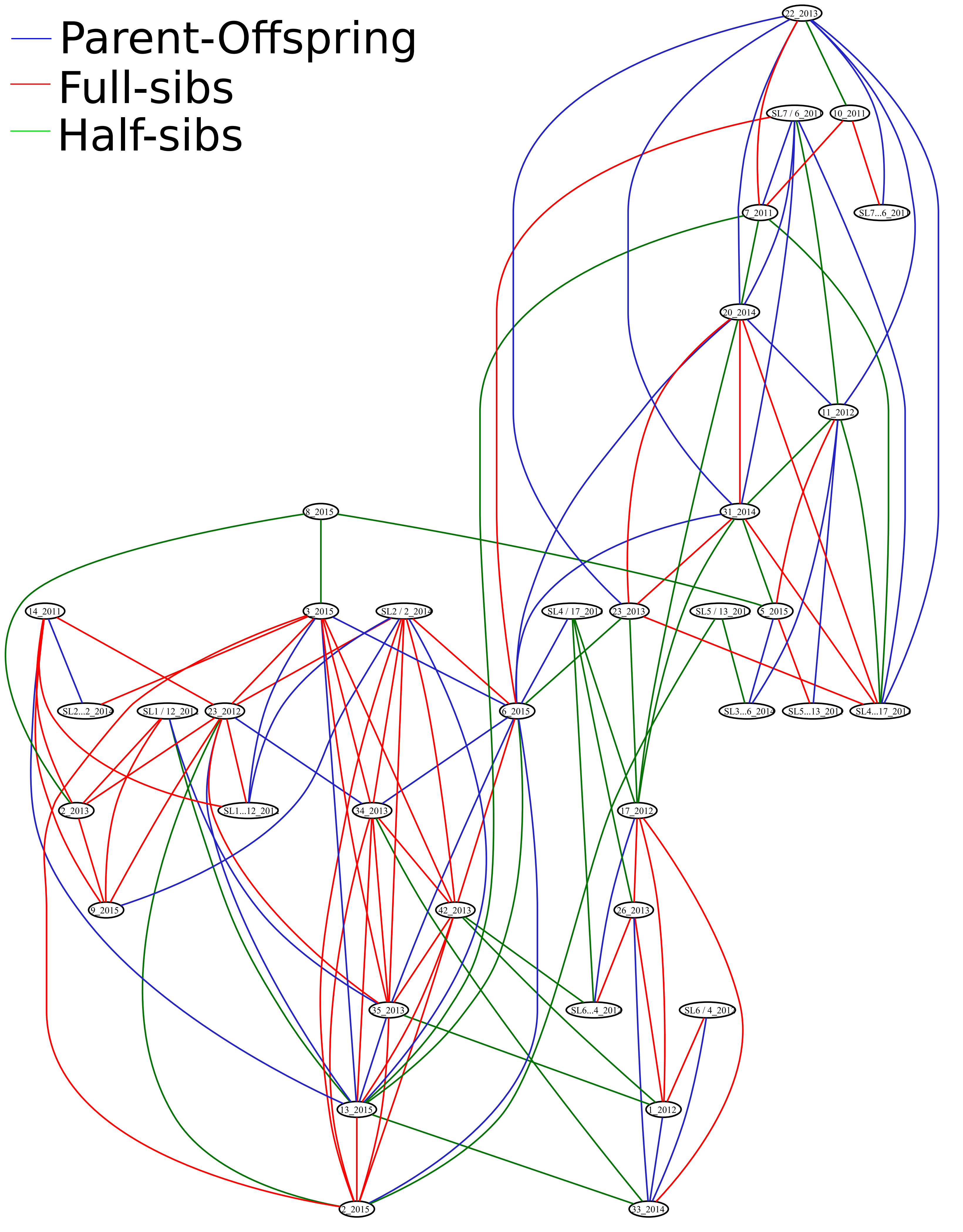
