## Supplemental Table 1 - Transects for "Diachronic monitoring of snow leopards at Sarychat-Ertash State Reserve (Kyrgyzstan) through scat genotyping: a pilot study" (from another version of this preprint: version 1, 2019-11-16)

**Table S1**: List of prospected transects from 2011 to 2015

| Index | Place | Date | Snow leopard evidences (0: no, 1: yes) | Number of scats harvested | Number of scats genotyped |
|---|---|---|---|---|---|
| W | After Kocheteuk | July 2014 | 0 | 0 | 0 |
| A | Atcha | June 2014 | 1 | 0 | 0 |
| | | June 2013 | 1 | 0 | 0 |
| P | Bir Baital | August 2015 | 1 | 1 | 0 |
| | | August 2014 | 1 | 1 | 1 |
| | | June 2014 | 1 | 3 | 3 |
| | | June 2013 | 1 | 3 | 2 |
| | | July 2012 | 1 | 1 | 1 |
| | | August 2012 | 1 | 2 | 2 |
| C | Bordu | June 2012 | 1 | 3 | 3 |
| | | June 2013 | 1 | 5 | 5 |
| B | Boroko | June 2013 | 1 | 0 | 0 |
| G | Chomoi | June 2013 | 1 | 6 | 6 |
| O | Eki Baital | August 2014 | 1 | 1 | 1 |
| | | July 2015 | 1 | 1 | 1 |
| | | August 2015 | 1 | 0 | 0 |
| Y | Near Bachkeul | August 2011 | 1 | 2 | 2 |
| | | June 2013 | 1 | 1 | 1 |
| F | Chong Bordu | June 2014 | 1 | 1 | 1 |
| I | Chong Koilou | June 2014 | 1 | 1 | 0 |
| M | Jaman-suu | June 2015 | 1 | 2 | 2 |
| | | July 2015 | 1 | 0 | 0 |
| | | August 2014 | 1 | 1 | 1 |
| | | August 2013 | 1 | 1 | 1 |
| | | August 2011 | 1 | 1 | 0 |
| R | Kichi Kashka tor | August 2015 | 0 | 0 | 0 |
| S | Chong Kashka tor | June 2014 | 1 | 1 | 1 |
| | | July 2015 | 0 | 0 | 0 |
| H | Kirk-choro | June 2014 | 1 | 6 | 6 |
| | | June 2013 | 1 | 5 | 5 |
| | | June 2012 | 1 | 0 | 0 |
| | | July 2012 | 0 | 0 | 0 |
| W | Kocheuteuk | July 2012 | 0 | 0 | 0 |
| | | August 2012 | 0 | 0 | 0 |
| | | August 2011 | 0 | 0 | 0 |

| | | | | | |
|---|---|---|---|---|---|
| | | July 2011 | 1 | 0 | 0 |
| Z | Koyendou | June 2015 | 0 | 0 | 0 |
| | | July 2015 | 0 | 0 | 0 |
| E | Orto-bordu | June 2013 | 1 | 3 | 3 |
| | | June 2012 | 1 | 0 | 0 |
| | | July 2012 | 1 | 1 | 0 |
| | | June 2014 | 1 | 1 | 1 |
| J | Orto Koilou | June 2014 | 1 | 1 | 0 |
| D | Kichi Bordu | June 2014 | 1 | 1 | 1 |
| K | Kichi Koilou | July 2012 | 0 | 0 | 0 |
| | | June 2013 | 0 | 0 | 0 |
| | | June 2014 | 1 | 1 | 0 |
| Q | Sary etchki | July 2015 | 0 | 0 | 0 |
| | | August 2014 | 0 | 0 | 0 |
| | | June 2013 | 1 | 5 | 4 |
| | | July 2011 | 1 | 5 | 5 |
| | | June 2012 | 1 | 3 | 3 |
| | | August 2011 | 1 | 0 | 0 |
| | | July 2012 | 1 | 1 | 1 |
| T | Sirdibai | July 2015 | 1 | 0 | 0 |
| | | June 2014 | 1 | 2 | 2 |
| | | August 2013 | 1 | 4 | 3 |
| | | June 2015 | 1 | 3 | 3 |
| | | July 2013 | 1 | 3 | 3 |
| | | August 2012 | 1 | 1 | 1 |
| | | July 2014 | 1 | 1 | 1 |
| | | August 2015 | 1 | 2 | 2 |
| | | July 2012 | 1 | 2 | 2 |
| | | July 2011 | 1 | 0 | 0 |
| | | August 2011 | 1 | 0 | 0 |
| | | June 2012 | 0 | 0 | 0 |
| U | Solomo | June 2014 | 1 | 3 | 3 |
| | | July 2014 | 1 | 1 | 1 |
| | | August 2014 | 1 | 2 | 2 |
| | | June 2015 | 1 | 2 | 2 |
| | | July 2015 | 1 | 1 | 1 |
| | | August 2015 | 0 | 0 | 0 |
| | | July 2013 | 1 | 4 | 4 |
| | | August 2013 | 1 | 1 | 0 |
| | | August 2012 | 0 | 0 | 0 |
| | | June 2012 | 1 | 3 | 3 |

| | | | | | |
|---|---|---|---|---|---|
| | | August 2011 | 1 | 2 | 2 |
| | | July 2011 | 1 | 3 | 3 |
| N | Uch Baital | July 2015 | 1 | 2 | 2 |
| | | August 2015 | 1 | 1 | 0 |
| | | July 2013 | 1 | 1 | 1 |
| | | August 2013 | 1 | 1 | 0 |
| | | June 2012 | 1 | 3 | 3 |
| | | July 2012 | 0 | 0 | 0 |
| | | August 2012 | 1 | 4 | 4 |
| | | July 2011 | 1 | 1 | 1 |
| | | August 2011 | 1 | 0 | 0 |
| V | Jili Boulak | June 2014 | 1 | 3 | 3 |
| L | Gueuleu | June 2014 | 1 | 13 | 13 |
| | | June 2013 | 1 | 7 | 7 |
| | | July 2012 | 1 | 1 | 1 |
